# Ecology and population genetics of the parasitoid *Phobocampe confusa* (Hymenoptera: Ichneumonidae) in relation to its hosts, *Aglais* species (Lepidoptera: Numphalidae)

**DOI:** 10.1101/2020.05.27.115212

**Authors:** Hélène Audusseau, Gaspard Baudrin, Mark R. Shaw, Naomi L.P. Keehnen, Reto Schmucki, Lise Dupont

**Affiliations:** Department of Zoology, Stockholm University, Stockholm, Sweden; UK Centre for Ecology & Hydrology, Wallingford, United Kingdom; Univ Paris-Est Creteil, CNRS, INRAE, IRD, IEES-Paris, F-94010 Creteil France; Sorbonne Université, IEES-Paris, F-75005 Paris, France; Université de Paris, IEES-Paris, F-75013 Paris, France; National Museums of Scotland, Chambers Street, Edinburgh, United Kingdom

**Keywords:** *A. urticae*, *A. io*, genetic variation, landscape heterogeneity, phenology

## Abstract

The biology of parasitoids in natural ecosystems remain very poorly studied, while they are key species for their functioning. Here we focused on *Phobocampe confusa*, a vanessines specialist, responsible for high mortality rates in very emblematic butterfly species in Europe (genus *Aglais*). We studied its ecology and genetic structure in connection with those of its host butterflies in Sweden. To this aim, we gathered data from 428 *P. confusa* individuals reared from 6094 butterfly larvae (of *A. urticae*, *A. io* and in two occasions of *Araschnia levana*) collected over two years (2017 and 2018) and 19 sites distributed along a 500 km latitudinal gradient. We found that *P. confusa* is widely distributed along the latitudinal gradient. Its distribution is constrained over time by the phenology of its hosts. The large variation in climatic conditions between sampling years explains the decrease in phenological overlap between *P. confusa* and its hosts in 2018 and the 33.5% decrease in the number of butterfly larvae infected. At least in this study, *P. confusa* seems to favour *A. urticae* as host: while it parasitized nests of *A. urticae* and *A. io* equally, the proportion of larvae is significantly higher for *A. urticae*. At the landscape scale, *P. confusa* is almost exclusively found in vegetated open land and near deciduous forests, whereas artificial habitats are negatively correlated with the likelihood of a nest to be parasitized. The genetic analyses on 89 adult *P. confusa* and 87 adult *A. urticae* using COI and AFLP markers reveal a low genetic diversity in *P. confusa* and a lack of population genetic structure in both species, at the scale of our sampling. Further genetic studies using high-resolution genomics tools will be required to better understand the population genetic structure of *P. confusa*, its biotic interactions with its hosts, and ultimately the stability and the functioning of natural ecosystems.

## 1. Introduction

Most biological studies of parasitoids have been done in the context of biocontrol in agricultural ecosystems. Such focus on parasitoids specialized on pest species, however, has limited our knowledge on the biology and function of parasitoids in natural ecosystems. For example, only a few of the over 100 000 ichneumonid species estimated are identified to date [1] and the biology and ecology of the vast majority of these species remain poorly understood [2–4]. Thus, while parasitoids constitute a large part of the biodiversity and are key species in the functioning of ecosystems, they have been widely neglected in ecological studies [3–7].

It is generally accepted that parasitoid species are sensitive to the interactions and population dynamics of their hosts and have their own habitat requirements [4]. However, little empirical evidence exists to adequately inform these processes and our knowledge of the ecology of most parasitoids is often based on sparse data obtained from a few randomly captured specimens [4]. The lack of data sets derived from systematic sampling limits our understanding of their distribution, in space and time, as well as the processes that drive their dynamics. Both the dynamics and the distribution of parasitoids are expected to be conditioned to that of their hosts [8]. This is also supported by studies of population genetic structure showing that parasitoids can track, locate and shift to different hosts in fragmented landscapes (reviewed in [9]). Comparing the spatial genetic structure of parasitoids with that of their hosts is a powerful approach that can provide essential understanding of species’ ecology and biotic interactions. The occurrence and survival of parasitoid populations also depend on a set of features of the habitat [10]. For example, the presence of sources of sugar and proteins during the reproductive season and appropriate shelters for overwintering are good indicators of habitat suitability for parasitoids (reviewed in [4,11]). At the landscape scale, however, the persistence of the parasitoid species is also likely to depend on their capability to disperse between suitable habitat patches. By affecting dispersal, habitat fragmentation and homogenization can have a negative impact on the population dynamics of parasitoids [12–14], with larger effects for species with limited dispersal capability [15,16]. The impact of habitat fragmentation on parasitoids is further exacerbated by the fact that they often occur at low densities, in populations that are therefore more likely to be vulnerable to changes [14,10]. The persistence of a population at a site is therefore the result of the interplay between local habitat suitability, species’ capacity to disperse between patches and the distribution in time and space of its potential hosts in the landscape.

*Phobocampe confusa* is an important parasitoid of emblematic butterfly species in Europe (genus *Aglais*). In Sweden, *P. confusa* represents the second cause of larval mortality due to parasitism in *A. urticae* and *A. io* [17]. *P. confusa* is an ichneumonid of the Campopleginae subfamily. It is a solitary endoparasitic koinobiont, that is, the female lays an egg in the body of its host, which continues to function and feed until the parasitoid larva emerges, in this case before the pupation of its host. The parasitoid overwinters as a pharate adult in the cocoon [18]. As in Hymenoptera generally, the sex-determination system of the species is haplodiploid, that is, females develop from fertilized eggs and are diploid, while males develop from unfertilized eggs and are haploid. The species is known to be a partly plurivoltine vanessine specialist and to parasitize the butterflies *Aglais io, Aglais urticae, Araschnia levana, Nymphalis polychloros* and *Polygonia c-album* [18], most often the first two. Although its effect on the abundance and dynamics of its hosts can be noticeable, the biology of *P. confusa* has not yet been systematically studied.

Here, we studied the ecology of *P. confusa* and how it interacts with its host butterfly species. We aimed to (i) identify the temporal constraints imposed by the phenology of its main host species in Sweden, *A. urticae* and *A. io*, (ii) investigate preference of hosts, and (iii) better understand the distribution of this parasitoid species in the landscape. In addition, as the population dynamic of parasitoids are likely to be closely linked to that of their hosts, (iv) we characterized and contrasted the genetic structure of *P. confusa* with one of its main host, *A. urticae*, to explore the potential biotic constraint induced by the host on the parasitoid and its dispersal.

## 2. Materials and Methods

### 2.1. Host butterflies

*Phobocampe confusa* has been recorded to parasitize several vanessine Nymphalidae species but in the vast majority of the recorded cases, *P. confusa* emerged from two nettle-feeding butterfly species, *Aglais io* and *A.urticae. Aglais urticae* and *A. io* are widely distributed over most of Sweden. These species are closely related butterflies [19] and show similar ecology. They are batch-laying species of 200 to 300 eggs, with larvae gregarious during the first three instars of their development, which then progressively become solitary. In Sweden, populations of *A. urticae* are partly bivoltine, depending on the weather conditions, with larvae observed in the field from May to the end of August. Populations of *A. io* are univoltine in Sweden and their phenology is slightly more restricted than for *A. urticae*, with larvae observed from late May to early August. Both *Aglais* species overwinter as adults.

Another Nymphalidae species which has recently established in the southern half of Sweden [20], *A. levana*, has occasionally been reported to be parasitized by *P. confusa*. Its spatial and temporal distribution overlaps greatly with that of *A. urticae* and *A. io.* Like *A. urticae* and *A. io*, it almost exclusively feeds on nettle (*Urticae dioica*). The species is also batch-laying, but with comparatively reduced batch size of 10 to 40 eggs. It is an obligate bivoltine species, with larvae observed in the field from June to early September. *A. levana* overwinters in the pupal stage.

### 2.2. Study area and data collection

Here we exploit the data collected in a large-scale field study of larval parasitism of nettle-feeding butterflies and described in Audusseau et al. [17]. In brief, the data correspond to the collection of larvae of four nettle-feeding butterflies occurring in Sweden, *A. io, A. urticae, A. levana*, and *V. atalanta*, over two years (2017-2018) and 19 sites along a 500 km latitudinal gradient in Sweden (Figure 1). The sites were selected to overlap, in comparable proportions among counties, habitats dominated by either agriculture lands or forests. At each site, we sampled nests of larvae fortnightly throughout the breeding season of the four butterfly species (early May to late August), for a total of 9 samplings per site. To maximize the diversity of the parasitoid species captured, we stratified the sampling design according to the developmental stage of the larvae (larval instars collected from 2^nd^ to 5^th^). This stratification enabled us to examine *P. confusa*’s attack preference for specific larval stages, as well as their time window of attack (see Material & Method in Audusseau et al. [17]). The development of butterfly larvae and the eventual emergence of parasitoids were monitored under controlled laboratory conditions. For the parasitized butterfly larvae, we recorded the larval stage and date at which parasitoids emerged from their cocoon. After emergence, freshly dead adult parasitoids were transferred to 95% alcohol to preserve the DNA for subsequent genetic analysis. For this study, we focused on the data on nests of larvae of *A. io, A. urticae*, and *A. levana*, and excluded data on *V. atalanta* as the species was not found to be parasitized by *P. confusa* [17]. For more details on the sampling protocol, sample size, winter diapause conditions, the complex of parasitoids and their relative distribution and abundance, see Audusseau et al. [17].

**Figure 1.**
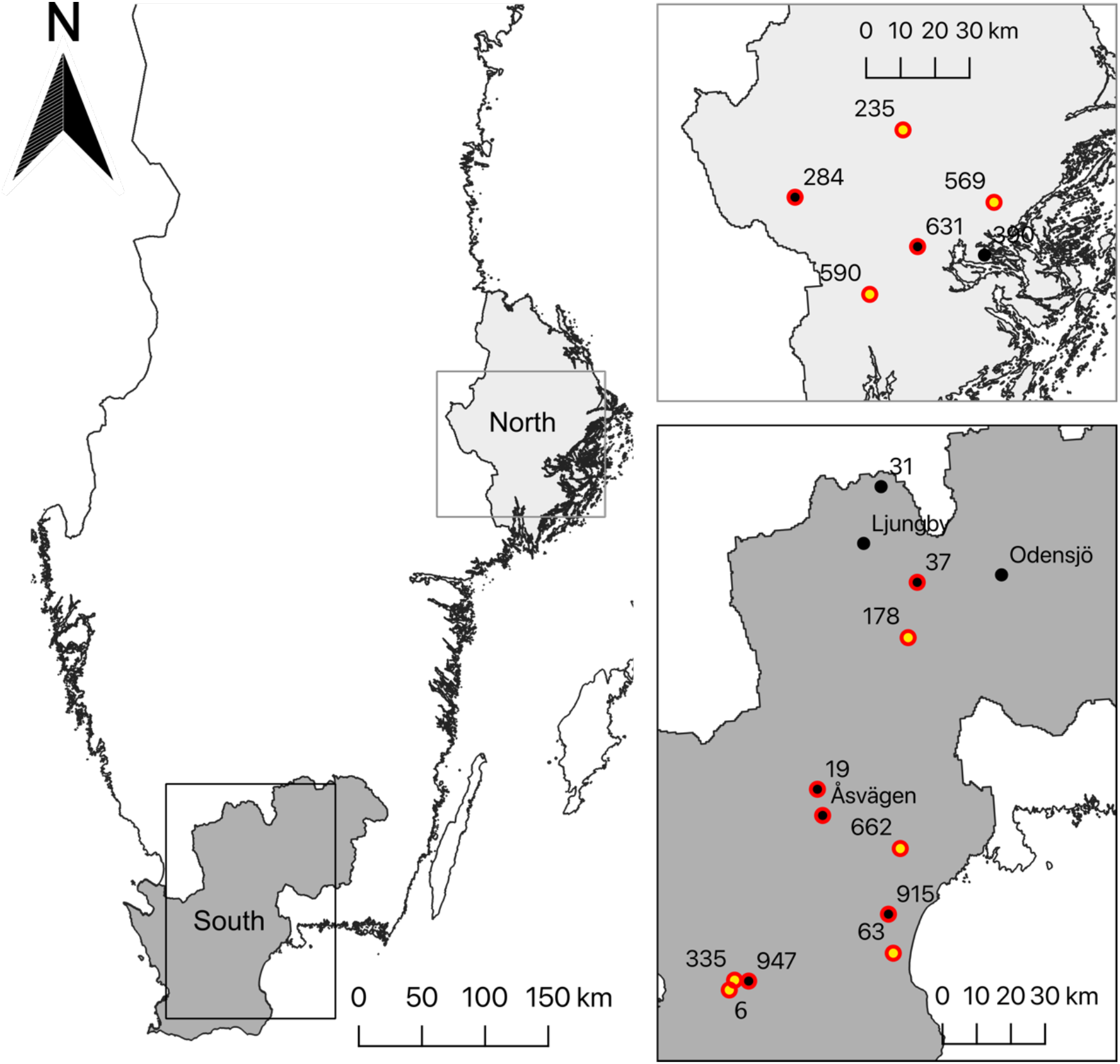
Map showing the 19 sites visited every two weeks during the two field campaigns (2017 and 2018). The sites are grouped into two regions, southern Sweden and the Stockholm region to the north. The points represent the location of the 19 sampled sites. The dots circled in red and the dots in yellow correspond to the sites where, respectively, individuals of *P. confusa* and *A. urticae* were used for genetic analyses.

### 2.3. Phenological synchrony between P. confusa and its hosts

We studied the temporal co-occurrence between *P. confusa* and *A. urticae, A. io,* and *A. levana*; that is, the phenological overlap between the parasitoid and its hosts. Specifically, we investigated differences in overlap between butterfly hosts and regions (south versus north) and controlled for differences between years. The phenological overlap was modelled using a linear model. The initial model included all the two-way interactions and model selection followed a backward elimination procedure.

The phenological overlap between *P. confusa* and its three host butterflies, or Overlap Parasitoid-Host index (OPH), was measured at each site *j* as the sum over the sampling weeks *k(1,…, 9)* of the minimum between the standardized abundance values of *P. confusa* (P*j,k*) and each of its hosts (H*j,k*) (eq. 1). For *P. confusa*, standardized abundance data (P*j,k*) refers to the number of individuals (NP) collected for a given sampling week *k* and site *j* and expressed in proportion of the total number of individuals of that species collected on all the samplings at the site *j* (eq. 2). For the host butterfly species (*A. urticae* or *A. io*), standardized abundance data refers to the number of nests collected for a given sampling week *k* and site *j* and expressed in proportion of the total number of nests of that species (NH) collected on all the samplings on the site *j* (eq. 3). The overlap index (OPH) is a parsimonious measure of the phenological overlap under the hypothesis that the parasitoid does not benefit from a surplus of resources [21]. The phenological overlap between species is calculated only when the two species, namely *P. confusa* and each of its hosts, were sampled at a site within a given year.

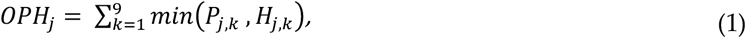

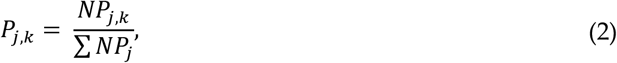

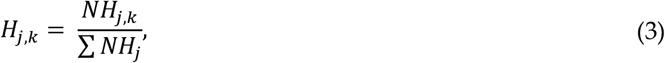

### 2.4. Pattern of attack

We investigated differences in *P. confusa* attack rates on its two main host butterflies, *A. urticae* and *A. io*, in two ways. First, we studied the proportion of butterfly nests parasitized by *P. confusa*. This analysis was restricted to butterfly nests sampled within the temporal window of occurrence of *P. confusa* (see Table S1) and at sites where *P. confusa* was observed at least once in the season (*n* = 359 nests). Second, we examined the proportion of larvae parasitized for each nest parasitized by *P. confusa* (*n* = 145 nests). The proportion of butterfly nests parasitized and the proportion of larvae parasitized by *P. confusa* per nest were modelled with a binomial error distribution.

We analysed variations in parasitism rates according to butterfly host, region, larval instar at collection, the phenological overlap, the year and week of collection, and the total number of butterfly nests of both host butterflies (*A. io* and *A. urticae*) occurring in the week of sampling. Based on preliminary exploration of the data, we included a quadratic term for the sampling week and phenological overlap. We also included the two-way interactions between the butterfly host and the region, the year, the larval instar at collection, the total number of butterfly nests at sampling, and the two-way interaction between region and year. Because few nests were collected at 1^st^ instar, we pooled them with nests collected at 2^nd^ instar. Model selection followed a backward elimination procedure. Model diagnostics were assessed using the R package DHARMa [22].

### 2.5. Habitat

We examined how habitat heterogeneity and fragmentation influenced the distribution of *P. confusa*. Using the models selected in the analyses of the proportion of butterfly nest parasitized and the proportion of larvae parasitized by *P. confusa* per nest (*see above*), we estimated the additional variance explained when including land cover variables. In the analyses, absences were informed by including data on butterfly nests collected at sites where *P. confusa* was not observed (*n* = 31), but that were sampled during its period of activity (Table S1). Land cover heterogeneity was modelled as the percentage of arable land, vegetated open land (e.g. field, meadow, grassland, offering easy running), deciduous forests, and artificial surfaces (buildings and roads) within the vicinity of the nests sampled. Habitat fragmentation was estimated from the total length of the edges measured between habitat types in the landscape surrounding each sampled nest. Land use heterogeneity and fragmentation were extracted from a land cover map produced at 10 m resolution by Naturvårdsverket (https://www.naturvardsverket.se/). To assess the effect of land cover on the propensity and intensity of parasitism, we computed each metric within buffers of increasing radius (10, 20, 30, 40, 70, 100, 200 and 500 meters) around each sampled nest. All metrics were calculated with the R packages sf [23] and raster [24]. The land cover classification of the Naturvårdsverket map followed the CORINE Land Cover level 3 (EEA, 2019). In our models, the proportion of butterfly nests parasitized and the proportion of larvae parasitized by *P. confusa* per nest were modelled with a binomial error distribution. Model selection followed a backward elimination procedure and models fit were assessed using the R package DHARMa [22].

### 2.6. Genetic structure of P. confusa and of A. urticae

The genetic structure of Swedish *P. confusa* and *A. urticae* were studied using two types of molecular markers, a fragment of the cytochrome c oxidase subunit (COI) mitochondrial gene and Amplified Fragment Length Polymorphism (AFLP). AFLPs have been commonly used to study the population genetic structure of species since the publication of the method by Vos et al. [25]. Although these dominant markers (defined by presence/absence) are less informative than Single Sequence Repeats (SSRs) or Single-Nucleotide Polymorphism (SNPs), AFLPs are more time efficient and less expensive, which make them suitable to study non-model species such as the ones examined here. Comparative studies have also shown that the genetic diversity found by SSRs and AFLPs are comparable, as the distribution over the entire nuclear genome of the latter counterbalances the performance of using a limited number of SSRs (<20 SSRs, [26]).

#### 2.6.1. DNA extraction

DNA was extracted from whole body tissue of 89 adult *P. confusa* collected across 15 sites, and from abdomenal material of 87 adult *A. urticae* (one butterfly individual per nest) collected across 8 sites spread across the latitudinal gradient using the NucleoSpin® 96 Tissue kit (Macherey-Nagel) (Figure 1). After extraction, the DNA samples were quantified and assessed using a spectrophotometer (NanoDrop® ND-1000 UV-Vis*; Thermo Scientific*) and we measured concentrations of about 30 ng/μL.

#### 2.6.2. Mitochondrial genetic variation

We sequenced the fragment of the COI gene proposed as a standard DNA barcode for animals [27] using LCO1490F and HCO2192R primers [28]. DNA sequencing was performed in both directions by Eurofins Genomics company and sequences were manually aligned using the BioEdit program. We estimated the diversity of haplotype and nucleotide using DNAsp v.5. software [29]. Afterwards, the relationships among haplotypes were examined using a haplotypic network constructed by a reduced-median algorithm [30] as implemented in the software NETWORK 4.1.1.1 (https://www.fluxus-engineering.com/sharenet.htm). We used a maximum parsimony algorithm to infer the most parsimonious branch connections between the haplotypes.

#### 2.6.3. Nuclear genetic variation

To study the nuclear genetic variation of *P. confusa*, only diploid females were used. Male Hymenoptera are haploids and carry only half of the genetic information that diploid females do. For this reason, using a mixture of both males and females could lead to ambiguous results. In addition, we genotyped only one individual per butterfly nest sampled in order to avoid genotyping of related individuals which would, potentially, reduce the genetic variability of our sample. We kept only non-ambiguous AFLP results, which led to a total of 39 *P. confusa* AFLP genotypes and 86 *A. urticae* AFLP genotypes.

We obtained the AFLP fragments from 600 ng of genomic DNA, digested successively with the TaqI and EcoRI restriction enzymes (1 h 30 at 65 and 37 °C., respectively for each enzyme). The digested DNA was incubated at 37 °C for 3 h in the presence of adapter pairs corresponding to both types of restriction sites and T4 DNA ligase enzyme (EcoRI top: 5’-CTCGTAGACTGCGTACC; EcoRI bottom: 5-AATTGGTACGCAGTCTAC; TaqI top: 5’-GACGATGAGTCCTGAC; TaqI bottom 5’-CGGTCAGGACTCAT) before amplifying them by two successive PCRs using the EcoRI-A and TaqI-A primers, during the pre-selective PCR, and TaqI-AAC and EcoRI-AAC (FAM) primers, during selective PCR. The separation of the labelled AFLP fragments and the acquisition of the raw fluorescence data was performed by the “Genomics” platform of the Henri Mondor Institute by capillary electrophoresis (Applied Biosystem) in the presence of the LIZ 500 size marker. The obtained AFLP profiles were calibrated and analysed using the GeneMapper© software (Applied Biosystems). Eight individuals of *P. confusa*, and 12 individuals of *A. urticae* were genotyped twice to estimate the genotyping error rate. AFLP genotyping followed the protocol described elsewhere [31–33].

The genetic diversity statistics, i.e. proportion of variable markers and gene diversity based on Nei’s formula [34], were calculated using AFLPdat program [35]. The spatial genetic structure for each of the two species were assessed by Bayesian inference, taking into account the multilocus AFLP genotype and the geographical coordinates of each individual [36], using the R package Geneland [37]. Individuals were grouped into genetic clusters representing homogeneous gene pools without a priori information about individual origin. We ran 5 replicate runs, with the number of clusters, K, ranging from 1 to 15, of a model of correlated frequencies, i.e. taking into account the similarity of the frequency of alleles between populations. We ran 100,000 iterations and sampled every 100 iterations.

## 3. Results

### 3.1. Patterns of occurrence of P. confusa

A total of 428 *P. confusa* individuals emerged from larvae collected from 146 different butterfly nests (Table 1), 257 in 2017 and 171 in 2018. *Phobocampe confusa* is the second most common parasitoid species found within our samples, beside *Pelatachina tibialis*, a weakly gregarious tachinid parasitoid of which we reared 1227 individuals out of the 526 butterfly larvae infested, collected from 165 different nests.

**Table 1.** Showing by region, year and butterfly host, and in order, in black the number of individuals of *P. confusa* reared and the number of butterfly nests parasitized by *P. confusa*, and in grey the total number of butterfly host larvae and the number of nests collected. Note that *A. levana* is not yet present in the north.

| year | Region/host | *A. urticae* | *A. io* | *A. levana* | Total by region |
| --- | --- | --- | --- | --- | --- |
| 2017 | North | 65/22/374/57 | 81/30/589/70 | - | 146/52/963/127 |
|  | South | 82/34/612/68 | 27/10/605/45 | 2/1/712/69 | 111/45/1929/182 |
|  | Total by species | 147/56/986/125 | 108/40/1194/115 | 2/1/712/69 | 257/97/2892/309 |
| 2018 | North | 6/4/598/58 | 11/3/379/26 | - | 17/7/977/84 |
|  | South | 78/23/669/66 | 76/19/685/63 | 0/0/871/98 | 154/42/2225/227 |
|  | Total by species | 84/27/1267/124 | 87/22/1064/89 | 0/0/871/98 | 171/49/3202/311 |

*Phobocampe confusa* was observed throughout the southern and northern regions of Sweden in both years, but its abundance in our samples varied between hosts, sites, counties and years (Table 1). Across sites and years, the abundance of *P. confusa* varied from 1 to 59 individuals per site in 2017 (18.21 ± 3.56, mean ± se) and from 1 to 82 in 2018 (13.15 ± 6.02, mean ± se). The species was absent from two sites in both years, site 31 and Odensjö. Additionally, *P. confusa* was not present in Ljungby and site 915 in 2017, and in 2018 it was absent from the sites 284, 569, 63, and Åsvägen. In our laboratory conditions, *P. confusa* adult emergence rate was of 29.0% with a total of 124 individuals that emerged, 48 males, 72 females and 4 that we failed to sex. All the emergence of adults of *P. confusa* occurred within the year of its cocoon formation. The low rate of emergence after winter diapause is probably the result of suboptimal husbandry of wintering cocoons.

*Phobocampe confusa* is a solitary parasitoid, laying one egg per larval host in most cases. Nevertheless, we observed one case where a larva of *A. io* was parasitized by both *P. confusa* and *Blondelia nigripes. Aglais urticae* and *A. io* are the two main hosts of *P. confusa* among the four butterfly species we sampled. 231 *P. confusa* larvae egressed from the 2254 *A. urticae* larvae collected, 196 out of the 2259 *A. io,* and 2 out of the 1583 *A. levana.*

### 3.2. Phenological synchrony between P. confusa and its hosts

The phenological overlap between *P. confusa* and its hosts varies significantly between year, butterfly hosts and region (Figure 2a, Table 2). While the phenological overlap between *P. confusa* and *A. urticae* and *A. io* are comparable in the north (estimate = −0.11 ± 0.11, t = −0.94, p = 0.35), the overlap is higher with *A. urticae* than with *A. io* in the southern region (estimate = 0.388 ± 0.147, t = 2.65, p = 0.010), and this for both years. Although we only recorded two cases where *P. confusa* parasitized *A. levana*, the phenological overlap between *P. confusa* and its host *A. levana* is comparable to the overlap observed for the native species *A. io* (estimate = −0.086 ± 0.090, t = −0.95, p = 0.34, Figure 2).

**Table 2.** Type II Anova table showing variation in phenological overlap between *P. confusa* and its hosts according to host species, region (south and north), year, and the two-way interaction between region and host species. R^2^_adj_ = 24.9, p < 0.001.

| Variables | Sum sq | Df | F | p |
| --- | --- | --- | --- | --- |
| Host | 0.802 | 2 | 6.17 | **0.004** |
| Region | 0.003 | 1 | 0.038 | 0.85 |
| Year | 0.347 | 1 | 5.34 | **0.024** |
| Region x host | 0.458 | 1 | 7.04 | **0.010** |
| Residuals | 4.031 | 62 |  |  |

**Figure 2.**
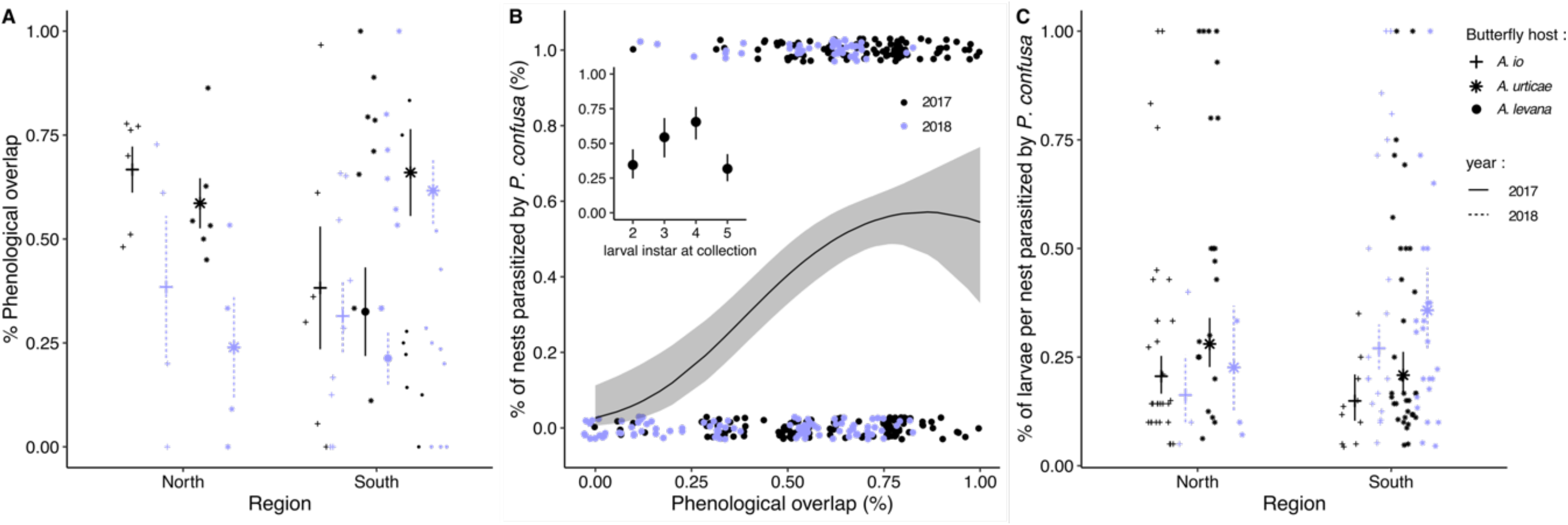
Plot showing (A) the phenological overlap between *P. confusa* and its hosts butterflies, *A. urticae*, *A. io* and *A. levana*, according to year and region; (B) the proportion of nests parasitized according to the phenological overlap and larval instar at collection; (C) the proportion of larvae parasitized by *P. confusa* per nest for *A. urticae* and *A. io* according to year and region. Dots represent the raw data, means ± confidence intervals. In purple are the data for 2018, in black for 2017. The shape of the dots refer to butterfly host species.

We also observe a significant decrease in the phenological overlap between *P. confusa* and its hosts in 2018 compared to 2017 (estimate = −0.144 ± 0.063, t = −2.31, p = 0.024, Figure 2a, Table 2). This probably reflects the considerable difference in temperature profiles between the two sampling years (Figure S1). In fact, if we replace the year variable by the corresponding cumulative growing degree-days above 13°C from January 1^st^ to August 31^st^ (GDD13), model selection procedure results in the same best model (SM 1). In contrast, precipitation from September to August (cumulative precipitation) is excluded from the final model, although it varied significantly between 2017 and 2018 (SM1).

### 3.3. Pattern of attack

The proportion of butterfly nests parasitized by *P. confusa* significantly varies with the larval instar at collection and shows a concave relationship with the phenological overlap (Table 3, Figure 2b). The proportion of butterfly nests parasitized by *P. confusa* increases with increasing phenological overlap and is higher for larval nests collected at the 3th and 4^th^ instar than for larvae collected at the 1^st^ and 2^nd^ instar and 5^th^ instar (Figure 2b). While the proportion of butterfly nests parasitized by *P. confusa* do not vary between the butterfly hosts, the proportion of larvae parasitized by *P. confusa* per nest is higher for *A. urticae* nests than for *A. io* nests (estimate = 0.41 ± 0.18, z = 2.27, p = 0.024, Figure 2c). The proportion of larvae parasitized by *P. confusa* also varies significantly between sampling years and this effect is specific to region. While in the northern region, the proportion of larvae parasitized by *P. confusa* per nest decreases between 2017 and 2018, the opposite is observed in the southern region (estimate = 1.04 ± 0.32, z = 3.24, p = 0.001). The proportion of larvae parasitized by *P. confusa* per nest also varies with the larval instar at collection (Table 3) and shows a concave relationship with the phenological overlap and the week of sampling (estimate phenological overlap^2^ = −4.47 ± 1.47, z = −3.04, p = 0.002; estimate sampling week^2^ = −0.09 ± 0.04, z = −2.61, p = 0.009, table 3, Figure 2c).

**Table 3.** Type II Anova table showing variation in parasitism rate according to the butterfly host, region, phenological overlap, larval instar at collection and the two-way interactions between the butterfly host and region, phenological overlap, and larval instar at collection.

| Variables | Proportion of nest parasitized |  |  | Proportion of larvae parasitism per nest |  |  |
| --- | --- | --- | --- | --- | --- | --- |
|  | LR Chisq | Df | *p* | LR Chisq | Df | *p* |
| Phenological overlap | 14.40 | 1 | < 0.001 | 10.40 | 1 | 0.001 |
| Phenological overlap<sup>2</sup> | 5.57 | 1 | 0.018 | 9.83 | 1 | 0.002 |
| Instar at collection | 25.78 | 5 | < 0.001 | 8.38 | 3 | 0.039 |
| Butterfly species | - | - | - | 5.08 | 1 | 0.024 |
| Year | - | - | - | 5.25 | 1 | 0.022 |
| Week of sampling | - | - | - | 7.90 | 1 | 0.005 |
| Week of sampling<sup>2</sup> | - | - | - | 7.36 | 1 | 0.007 |
| Region | - | - | - | 0.47 | 1 | 0.49 |
| Region x Year | - | - | - | 11.57 | 1 | < 0.001 |

### 3.4. Habitat

The effect of land cover heterogeneity and fragmentation is relatively constant between 10 to 200m radius around the butterfly nests sampled, and is not detected at 500m radius, possibly due to the overlap in landscape buffers around each butterfly nest at that scale. For this reason, we focus on the results for the effect of land cover within a 100m buffer radius and present the details of the models for each buffer zone as supplementary material (Table S2 & S3 in SM3). We find that the likelihood of a butterfly nest to be parasitized by *P. confusa* decreases with increasing proportion of artificial surface (estimate artificial surface 100m = −0.0467 ± 0.022, z = −2.17, p = 0.030, Figure 3), whereas the proportion of larvae parasitized by nest increases (estimate artificial surface 100m = 0.026 ± 0.011, z = 2.54, p = 0.024, Figure 3). We also observe a positive effect of the proportion of deciduous forest in the vicinity of the nest on the proportion of larvae parasitized by nest (estimate deciduous 100m = 0.018 ± 0.005, z = 3.39, p < 0.001, Figure 3).

**Figure 3.**
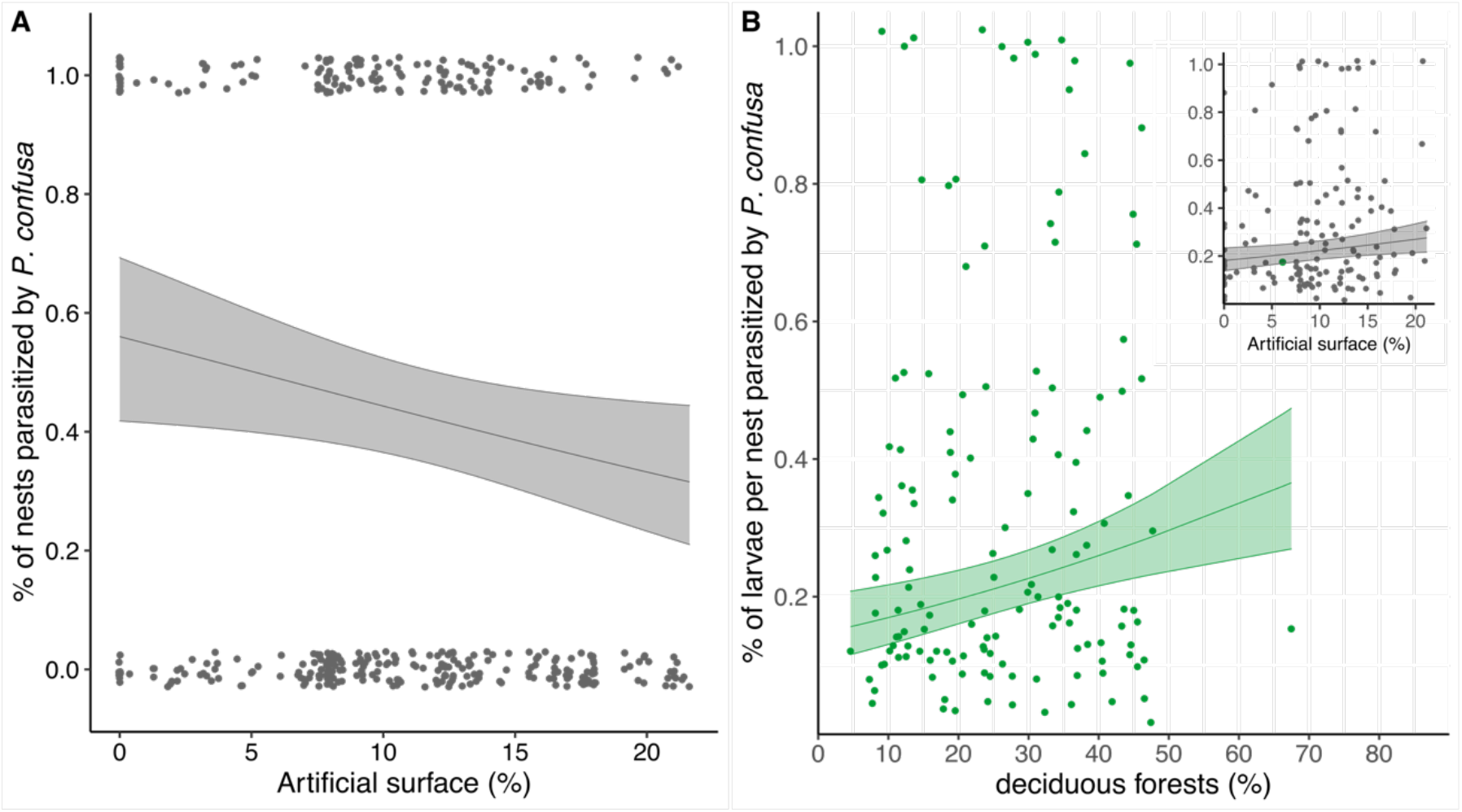
Plot showing (A) the proportion of nests parasitized according to the proportion of artificial surface within a buffer zone of 100m radius and (B) the proportion of larvae parasitized by *P. confusa* per nest according to the proportion of deciduous forests and artificial surface within a buffer zone of 100m radius. Dots correspond to the raw data, means ± confidence intervals correspond to the estimated marginal means from the model.

Note that this analysis focuses on the impact of land cover types well represented in the vicinity of the nests sampled, which are arable land, vegetated open land (e.g. field, meadow and grassland), deciduous forests, and artificial surfaces (building and road) (Figure S2). Although we initially selected sampling sites in landscapes (1km radius) with diverse land covers, butterfly nests were located (within 10m) in 87.4% of the cases near open vegetated land and in 58.5% of the cases near deciduous forests, stressing the importance of these two land covers for the species (Figure S2).

### 3.4. Genetic structure of P. confusa and of A. urticae

#### 3.4.1. Mitochondrial and nuclear genetic variation of *P. confusa*

For *P. confusa*, we obtained 88 sequences of a 613 bp fragment of the COI gene (GenBank Accession Numbers). We detect 5 haplotypes (Figure 4, Table 4) defined by 2 parsimony informative sites, among 4 variable sites. The global haplotype diversity and nucleotide diversity are of 0.284 and 0.00051 respectively. Over the 82 AFLPs fragments recorded, only 15 are polymorphic, for which no error of genotyping was observed in replicates. We observe extremely low genetic diversity indices in the North and South regions (Table 4). Bayesian inference revealed no genetic structuring and only one genetic cluster was identified by Geneland V 4.0.3 [37].

**Table 4.**
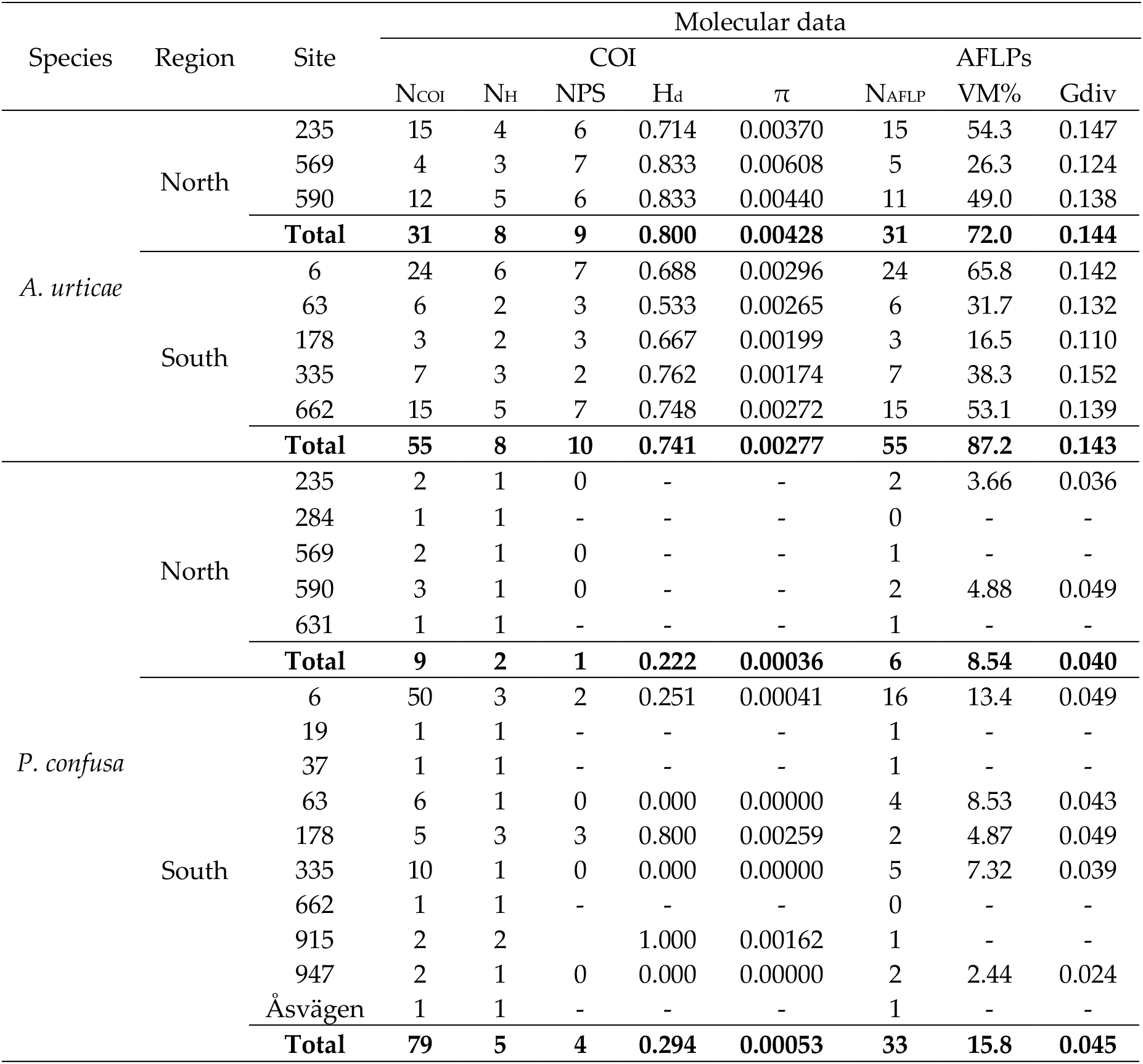
Genetic variation within *A. urticae* and *P. confusa* populations estimated using COI mitochondrial gene and AFLPs. Sample size (N_COI_ and N_AFLP_), number of COI haplotype (N_H_), number of polymorphic site (NPS), haplotype diversity (H_d_), nucleotide diversity (π), percentage of variable markers (VM%) and gene diversity (Gdiv).

| Species | Region | Site | Molecular data |  |  |  |  |  |  |  |
| --- | --- | --- | --- | --- | --- | --- | --- | --- | --- | --- |
|  |  |  | COI |  |  |  |  | AFLPs |  |  |
| | | | $N_{\text{COI}}$ | $N_{\text{H}}$ | NPS | $H_{\text{d}}$ | $\pi$ | $N_{\text{AFLP}}$ | VM% | Gdiv |
| *A. urticae* | North | 235 | 15 | 4 | 6 | 0.714 | 0.00370 | 15 | 54.3 | 0.147 |
|  |  | 569 | 4 | 3 | 7 | 0.833 | 0.00608 | 5 | 26.3 | 0.124 |
|  |  | 590 | 12 | 5 | 6 | 0.833 | 0.00440 | 11 | 49.0 | 0.138 |
|  |  | **Total** | **31** | **8** | **9** | **0.800** | **0.00428** | **31** | **72.0** | **0.144** |
|  | South | 6 | 24 | 6 | 7 | 0.688 | 0.00296 | 24 | 65.8 | 0.142 |
|  |  | 63 | 6 | 2 | 3 | 0.533 | 0.00265 | 6 | 31.7 | 0.132 |
|  |  | 178 | 3 | 2 | 3 | 0.667 | 0.00199 | 3 | 16.5 | 0.110 |
|  |  | 335 | 7 | 3 | 2 | 0.762 | 0.00174 | 7 | 38.3 | 0.152 |
|  |  | 662 | 15 | 5 | 7 | 0.748 | 0.00272 | 15 | 53.1 | 0.139 |
|  |  | **Total** | **55** | **8** | **10** | **0.741** | **0.00277** | **55** | **87.2** | **0.143** |
| *P. confusa* | North | 235 | 2 | 1 | 0 | - | - | 2 | 3.66 | 0.036 |
|  |  | 284 | 1 | 1 | - | - | - | 0 | - | - |
|  |  | 569 | 2 | 1 | 0 | - | - | 1 | - | - |
|  |  | 590 | 3 | 1 | 0 | - | - | 2 | 4.88 | 0.049 |
|  |  | 631 | 1 | 1 | - | - | - | 1 | - | - |
|  |  | **Total** | **9** | **2** | **1** | **0.222** | **0.00036** | **6** | **8.54** | **0.040** |
|  | South | 6 | 50 | 3 | 2 | 0.251 | 0.00041 | 16 | 13.4 | 0.049 |
|  |  | 19 | 1 | 1 | - | - | - | 1 | - | - |
|  |  | 37 | 1 | 1 | - | - | - | 1 | - | - |
|  |  | 63 | 6 | 1 | 0 | 0.000 | 0.00000 | 4 | 8.53 | 0.043 |
|  |  | 178 | 5 | 3 | 3 | 0.800 | 0.00259 | 2 | 4.87 | 0.049 |
|  |  | 335 | 10 | 1 | 0 | 0.000 | 0.00000 | 5 | 7.32 | 0.039 |
|  |  | 662 | 1 | 1 | - | - | - | 0 | - | - |
|  |  | 915 | 2 | 2 |  | 1.000 | 0.00162 | 1 | - | - |
|  |  | 947 | 2 | 1 | 0 | 0.000 | 0.00000 | 2 | 2.44 | 0.024 |
|  |  | Åsvägen | 1 | 1 | - | - | - | 1 | - | - |
|  |  | **Total** | **79** | **5** | **4** | **0.294** | **0.00053** | **33** | **15.8** | **0.045** |

**Figure 4.**
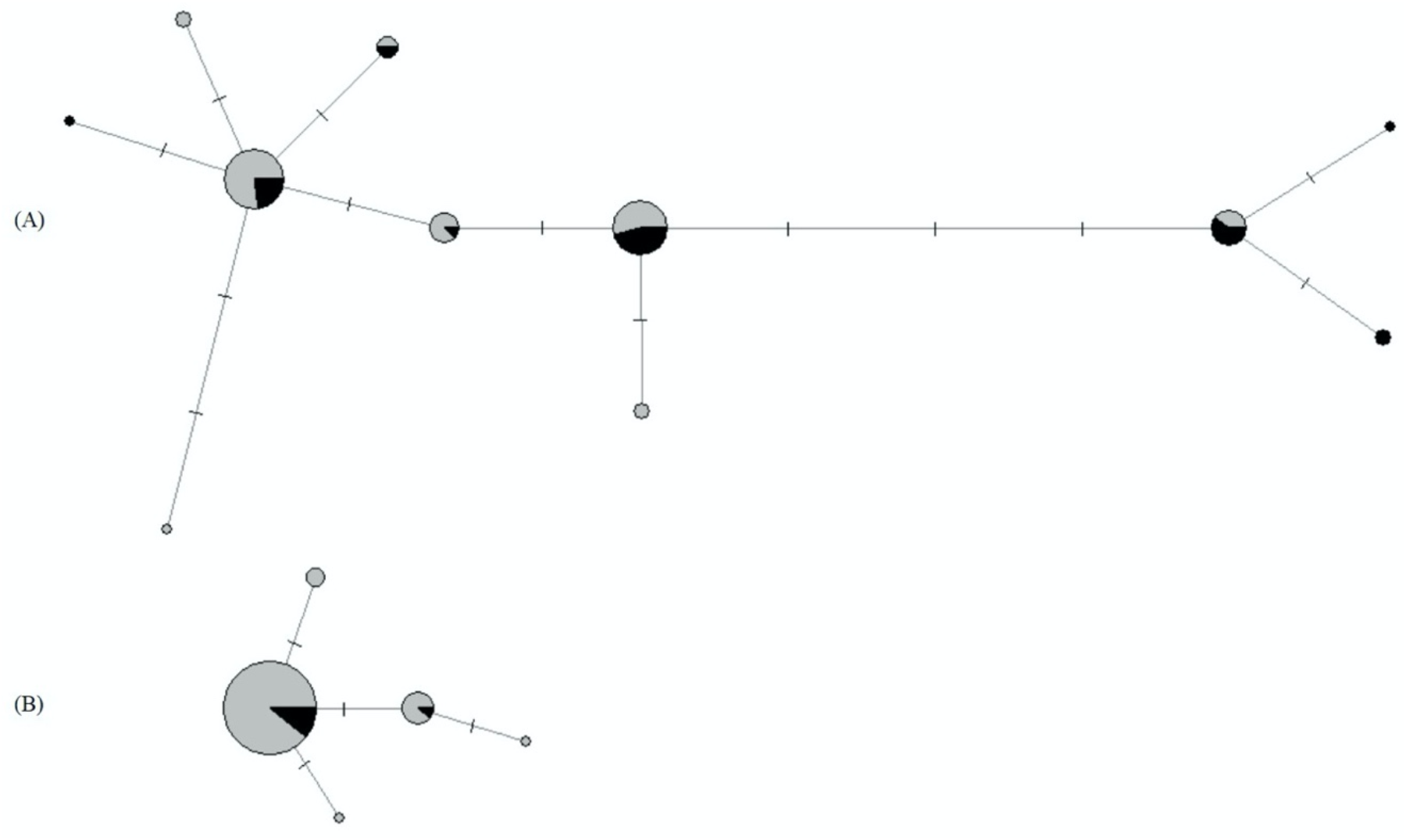
COI gene haplotype network for (A) *A. urticae* samples and (B) *P. confusa* samples. Circle size is relative to the proportion of each haplotype in the sample. Mutational steps are indicated by lines. Individuals collected in the South of Sweden are in grey, individuals collected in the North of Sweden are in black.

#### 3.4.2. Mitochondrial and nuclear genetic variation of *A. urticae*

For *A. urticae*, we obtained 86 sequences of a 603 bp fragment of the COI gene (GenBank Accession Numbers). We detect 11 haplotypes (Figure 4, Table 4) defined by 9 parsimony informative sites, among 13 variable sites. The global haplotype diversity and nucleotide diversity are of 0.775 and 0.00349 respectively. We obtained a total of 243 polymorphic AFLPs fragments with a very low genotyping error rate (< 1%). We do not observe a significant difference in gene diversity between regions (Table 4). In addition, the Bayesian inference did not show a genetic structuring of our data, only one genetic cluster was identified by Geneland V 4.0.3 [37].

## 4. Discussion

The total number of larvae infected by *P. confusa* has decreased by 33.5 % between 2017 and 2018. This is not related to a reduction in host availability as, with comparable sampling effort, between the two years the number of butterfly larvae collected increased by 6.9 % for the two long-native host butterflies and by 10.72% when including *A. levana.* The observed decrease is most likely explained by the very peculiar climatic conditions recorded in 2018 as that year was exceptionally dry in Scandinavia with both an increase in average temperature over the season and lower precipitation (see SM 1). In turn, the variation in climatic conditions explains a large part of the observed decreased in phenological overlap between *P. confusa* and its native hosts. This decrease was most pronounced in the northern region and resulted in the low number of reared *P. confusa.* There, the proportion of native butterfly nests parasitized by *P. confusa* dropped from 40.9 % in 2017 (52 out of 127 native butterfly nests sampled) to 8.05 % in 2018 (7 out of 84 native butterfly nests sampled). In addition to the importance of the overlap between the phenology of the host butterflies and *P. confusa*, the probability of detecting a case of a nest parasitized by *P. confusa* is strongly influenced by the larval stage at the time of collection and was highest for the nests for which the larvae were collected in the fourth larval instar. We estimate the time window of attack of a larval host by *P. confusa* to be of at least a week in the wild and probably longer for *A. io* than *A. urticae* due to its longer development time (see SM 2). We did not find any difference between native species in the probability of a nest to be parasitized; however the intensity of parasitism, taken as the proportion of larvae parasitized per nest, differs between species and is significantly higher for *A. urticae* than for *A. io.* This result suggests that, at least in this study, *P. confusa* seems to favour *A. urticae* as host.

The large between-year variation in climatic profile highlights the potential impact of warming on our study system. Climate change is a challenge for ectothermic species such as parasitoids and their butterfly hosts. As they do not produce heat, their development and survival rely on the temperature of their habitat [38]. In Sweden, and more generally at higher latitudes where the magnitude of the warming is greater [39], we stronger effects of climate change. In that respect, we found a negative impact of the modification of the climatic profile in Sweden on *P. confusa.* This aligns with previous studies showing that specialist species, as is the case for *P. confusa*, are particularly sensitive to climatic unpredictability [40,41]. However, this contrasts with the overall pattern of parasitism as Audusseau et al. [17] reported a higher level of parasitism (all parasitoid species combined) in 2018. Alternatively, at northern latitudes the impact of climate change is modulated by the fact that most species are living at much lower temperature than their physiological optima and, for those, warming is expected to enhance individual fitness [42]. Most importantly, climate warming may alter life history traits of both the parasitoids and their hosts [38,43,44], causing rapid mismatch in the phenology of these interacting species [45], as shown in our data. Host use might also be affected by the warming. In that respect, it is important to stress that *A. levana* has recently established in Sweden, probably as a result of climate warming [20]. Here, we only reported two cases of *A. levana* larvae parasitized by *P. confusa.* This low level of parasitism might be explained by the enemy release hypothesis [46,47], which predicts that when establishing in a new area, the species escape their natural enemies until the local parasite complex recolonizes the species. However, *A. levana* is a potential host for *P. confusa* and the phenologies of these two species greatly overlap in Sweden, suggesting that *A. levana* could provide a refuge for *P. confusa* at a time when the native hosts are rare. Future monitoring of parasitism in *A. levana* and comparative data on the attack rate by *P. confusa* on *A. levana* in other parts of the butterfly’s range, and where the species are known to co-occur, would be insightful in that respect.

We found that butterfly nests and, therefore *P. confusa*, preferentially occur in habitat characterized by vegetated open land and where deciduous forests are found in the close vicinity. At a scale of 10 m radius around the butterfly nests sampled, the surrounding habitat of 87.4% of the nests included open vegetated land and for 58.5% deciduous forests. Association with these habitats might partly be explained by the pattern of distribution of nettles, *Urtica dioica*, the (practically exclusive) host plants of these butterflies. Nettles, common in northern Europe, are found in a diverse range of habitats but preferentially in nutrient-rich soils and in sites with moderate shading [48]. They are also found in deciduous woodland when the earth soil properties and insolation conditions are sufficient [48], but our field experience in Sweden showed that butterfly nests are generally found on nettle stands located along field edges of cultivated land or roads, in grasslands, meadows, and grazed fields, habitats classified as open vegetated land in the CORINE Land Cover classification (level 3). While this suggests a reduced importance of deciduous forest, this habitat could play an important role and provide a good refuge for the species. This is supported by the observed increase in the proportion of larvae parasitized per nests in landscapes with higher proportion of deciduous forest. We further detected a significant impact of the proportion of artificial surface on the occurrence of *P. confusa.* The probability of a butterfly nest to be parasitized by *P. confusa* decreased significantly with increased proportion of artificial surface, but the proportion of larvae parasitized per nest significantly increases. Other studies have shown that parasitoids suffer from environmental changes such as habitat fragmentation and habitat loss (e.g. [14,12,49]). We did not detect a specific effect of fragmentation, but fragmentation is highly correlated with the proportion of artificial surface (within a buffer of 100m radius, R^2^ = 0.65, p < 0.001), which has a significant negative impact on the propensity of a nest to be parasitized. The mechanisms by which artificial surfaces influence the distribution of *P. confusa* are difficult to assess and would require further experiments. Among the potential explanations, the alteration (and unevenly) of the nutritional content of nettles at close proximity to human habitation, or habitat fragmentation, may alter the parasitoids searching behaviour and their ability to find a nettle patch and/or might be associated with a higher mortality during the overwintering period, weakening the local populations (reviewed in [50]). The position of parasitoids in the food chain further increases their vulnerability to environmental changes [3,5].

To date, no genetic data have been made available for *P. confusa*. Here, we show that the COI genetic diversity is extremely low in this species, at least within the geographical scale of our study. We found only five different haplotypes which diverged by no more than 3 mutational steps (Figure 4). The lack of variability, which was confirmed at the nuclear level using AFLPs data, could suggest a recent spread of bottlenecked populations or could be the result of inbreeding. Population genetic theory indeed demonstrates that inbreeding is possible in haplodiploids [51] because the purging of deleterious and lethal mutations through haploid males reduces inbreeding depression (i.e. the lower fitness of offspring of genetically related parents compared to that of unrelated parents [52]). Solitary haplodiploid species, such as *P. confusa*, are however assumed to be primarily outbred while gregarious haplodiploid wasps (i.e. those that deposit more than one egg per host) are more likely to have a history of inbreeding [53]. This lack of genetic variability made it impossible to discern a population structure for *P. confusa* at the geographical scale of our study. In comparison, the COI genetic diversity observed in our samples of *A. urticae* was higher, with a total of 11 haplotypes (Figure 4). Although an important number of polymorphic AFLPs fragments (243) were obtained in our dataset, the spatial genetic analysis did not reveal any population genetic structure in *A. urticae*. This result is in concordance with previous studies on *A. urticae*, wherein long-distance gene flow is suggested to be important in this species. Using allozyme loci, Vandewoestijne et al. [54] have suggested that the population genetic structure of *A. urticae* at a regional scale is characterized by high movement rates between neighbouring patches, long-distance migration and rare extinction/recolonization events. A more recent study using mitochondrial sequences of the cytochrome c oxidase I mitochondrial gene (COI) and the control region showed that at the scale of the distribution of the species high gene flow is the primary factor shaping its population genetic structure [55].

Further studies at a larger geographical scale are needed to fully understand the relationship between the population genetic structure of *P. confusa* and that of its host, since the dispersal ability of the host *A. urticae* is larger than the geographical scale investigated in this study. Although AFLPs was successfully used in this study (i.e. we obtained more than 250 polymorphic markers in the lepidopteran host), high-resolution genomics tools, such as restriction- site DNA sequencing (RADseq, [56]), could provide additional information. Here, we highlight that further genetic studies on *P. confusa* and on all its potential hosts are required to understand the pattern of distribution of the species in the landscape in relation to that of its hosts. This would also allow further investigations of the dispersal ability of this species, an essential component for conservation ecology perspectives.

## 5. Conclusions

In this study, we focused on a parasitic hymenopteran, which represents one of the most species-rich insect groups [57], to provide insights into the ecology and the genetics of *Phobocampe confusa,* in relation to the one of its host butterflies in Sweden. So far, our knowledge of the ecology of this parasitoid was mainly limited to the work from Pyörnilä [58] (in which *P. confusa* was misidentified as *Hyposoter horticola*), although the species causes high mortality rates in very common and emblematic butterfly species in Europe. In particular, we showed that the occurrence of *P. confusa* relies on its phenological match with its host butterflies. It attacks similarly nests of *A. urticae* and *A. io*; however the proportion of larvae parasitized per nest is higher for *A. urticae.* Within our sample, the species occurred preferentially in vegetated open land and showed a high dependence on the occurrence of deciduous forests in the near surrounding. Artificial surfaces, however, seem to have a negative impact on the distribution of *P. confusa*. The genetic analyses did not reveal a population genetic structure in our study population, and further work is required to understand what is structuring the population genetics of *P. confusa*, understand its dispersal abilities and its biotic interactions with its hosts. Such knowledge is crucial to further our understanding of the factors and mechanisms shaping the stability and the functioning of natural ecosystems, including for conservation efforts.

## Supplementary Materials

Supplementary Material 1: Climatic variations between years and counties; Supplementary Material 2: Phenology and temporal window of attack of the host by *P. confusa*; Supplementary Material 3: Habitat characteristics associated with *P. confusa* occurrence for buffer zone radii varying from 10 to 500 m.

## Author Contributions

Conceptualization, H.A., L.D. and R.S.; Data collection, H.A and N.K..; taxonomic identification, M.R.S; Ecological analyses H.A., G.B., and M.R.S.; Genetic Analyses, G.B. and L.D; Funding acquisition H.A. and L.D.; H.A. and L.D. have written the original draft and all the authors contributed substantially to the revisions; All authors have agreed to the published version of the manuscript.

## Acknowledgments

We acknowledge the E-OBS dataset from the EU-FP6 project UERRA (http://www.uerra.eu) and the Copernicus Climate Change Service, and the data providers in the ECA&D project (https://www.ecad.eu). We thank Saskya van Nouhuys for enlightening discussions and Maria de la Paz Celorio-Mancera and Nils Ryrholm for their assistance in this project.

