## Supplementary materials 1, 2 and 3 for "Ecology and population genetics of the parasitoid *Phobocampe confusa* (Hymenoptera: Ichneumonidae) in relation to its hosts, *Aglais* species (Lepidoptera: Numphalidae)"

**Supplementary Material 1: Climatic variations between years and counties (Sweden).**

Temperature data were acquired throughout the field sampling with tinytag recorders (Tinytag Plus 2, TGP-4020) placed in the field in nettle patches at a height between 50 and 80 cm. We tested for the differences between counties and years in growing degree day-base 13°C (GDD13) accumulated over the reproductive season (from earliest May 9th to latest August 29th). For that, we modelled GDD13 using a generalized additive model with a normal error distribution including year, county and the interaction between year and county as linear effects, and the Julian day as a non-linear effect. We observed significant differences in GDD13 accumulated over the reproductive season between counties ($F = 134.9$, $p < 0.001$) and years ($F = 2333.5$, $p < 0.001$, Fig. S1). Between years, the relative change in the GDD13 accumulated was highest in the counties of Skåne and Kronoberg (estimate = 131.2, $t = 48.3$, $p < 0.001$ and estimate = 17.4, $t = 5.1$, $p < 0.001$, in Kronoberg and Skåne, respectively) than in the county of Stockholm (estimate = -37.3, $t = 3.7$, $p < 0.001$) (Fig. S1). In Skåne, GDD13 at the end of the reproductive season was of 217.49 ± 14.41°C in 2017 and of 531.13 ± 35.28°C in 2018. In Kronoberg, GDD13 at the end of the reproductive season was of 155.24 ± 9.27 °C in 2017 and of 427.65 ± 35.83°C in 2018. In Stockholm, GDD13 at the end of the reproductive season was of 260.78 ± 24.96°C in 2017 and of 518.70 ± 30.91°C in 2018.

Precipitation data were extracted for each site from the E-OBS v19.0e ([1], https://www.ecad.eu/). The resolution of these data is of 0.1 degree, which is about 11.11 km. We extracted these data in R 3.6.1 [2], using the packages ncdf4, raster, rgdal, sf, and lubridate [3–7]. Here, we only considered precipitations during the reproductive season of our study species; that is, precipitation from May 1st to August 31st. We modelled cumulative precipitation using a generalized additive model with a normal error distribution including year, county and the interaction between year and county as linear effects, and the Julian day as a non-linear effect. The cumulative precipitation was log-transformed prior to inclusion in the model. We observed significant differences in cumulative precipitation over the reproductive season between counties ($F = 284.3$, $p < 0.001$) and years ($F = 7.8$, $p = 0.005$, Fig. S1). Between years, the relative change in cumulative precipitation was higher in the counties of Skåne and Stockholm (estimate = -0.91, $t = -23.29$, $p < 0.001$ and estimate = -0.42, $t = -10.52$, $p < 0.001$, in Skåne and Stockholm, respectively) than in the county of Kronoberg, which showed the least change in precipitation profile between years (estimate = -0.09, $t = -2.80$, $p = 0.005$) (Fig. S1).

Thus, associated with the overall increase in average temperature over the season, the precipitation was significantly lower in 2018 compared to 2017. 2018 was an abnormally dry year with respect to 1981-2010 (https://climate.copernicus.eu/dry-and-warm-spring-and-summer).

The between years variation in temperature profile is correlated to the observed change in phenological overlap between years. Following the same procedure as described in the manuscript section 2.3., and replacing the variable year by GDD13, model selection procedure resulted in the same best model (region: $F = 0.005$, $p = 0.94$; GDD13: $F = 4.75$, $p = 0.03$; host:

F = 6.19, p = 0.004; region x host: F = 6.96, p = 0.01). The explained variance of this model is comparable, of 24.2%.

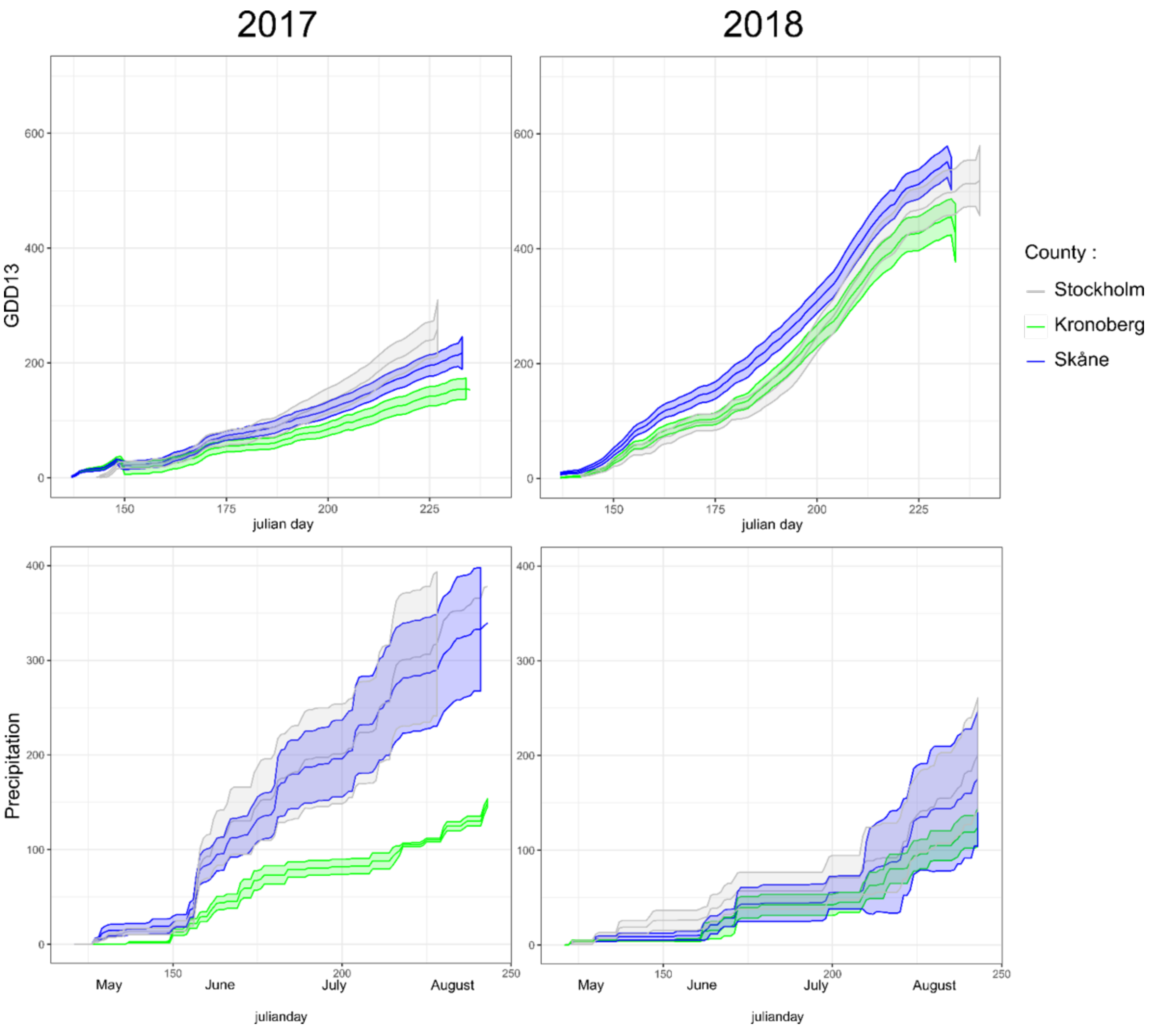


**Figure S1** Cumulative growing degrees-days above 13°C and precipitation in Kronoberg, Skåne, and Stockholm in 2017 and 2018.

**Acknowledgments:** We acknowledge the E-OBS dataset from the EU-FP6 project UERRA (http://www.uerra.eu) and the Copernicus Climate Change Service, and the data providers in the ECA&D project (https://www.ecad.eu). AH acknowledges support from the Swedish Research Council (2016-06737).

**Supplementary Material 2: Phenology and temporal window of attack of the hosts (*Aglais urticae* and *A. io*) by *Phobocamp. confusa.***

Parasitism by *Phobocampe confusa* started mid-May, both in the south and the north of Sweden and in both years of our study (2017 and 2018). In 2017, cases of parasitism by *P. confusa* was found until mid-July in the north and until the beginning of August in the south. The time window of the occurrence of *P. confusa* was substantially shorter in 2018 with the last occurrence of the species being recorded 4 and 6 weeks earlier in the northern and the southern regions, respectively (year = 1.81, t = -1.75, p = 0.092). The reduction in the time window of occurrence of *P. confusa* was most pronounced in the north (average time widow across sites in the north: 2017 = 6.0 ± 1.10 weeks, 2018 = 2.0 ± 1.15 weeks) compared to the south (average time widow across sites in the south: 2017 = 5.0 ± 3.70 weeks, 2018 = 4.33 ± 2.65 weeks), even though in both regions the difference between was not significant.

*Phobocampe confusa* emerged from *A. urticae* larvae collected from the 2 to 5$^{th}$ instar. We detected no evidence of *P. confusa* parasitism on first instar larvae of *A. urticae* (that is 51 larvae collected across 8 nests). The temporal window of attack of *P. confusa* for this host corresponds to the time during which *A. urticae* develop from 2$^{nd}$ to 4$^{th}$ instar, that is on average 5.15 days at 23°C and 22L:2D light regime (our laboratory rearing conditions).

*Phobocampe confusa* was found to emerge from *A. io* larvae collected from 2$^{nd}$ to 5th instar and from 5 larvae from one nest collected at the first instar *A. io* (that is 5 larvae out of 32 larvae collected from 1 out of the 6 nests of first instar larva sampled). The temporal window of attack of *P. confusa* for this host is on average of 7.80 days at 23°C and 22L:2D light regime.

Parasitism rate was highest when larvae were collected at the 4$^{th}$ instar. On the other hand, parasitism rate for larvae collected at the fifth instar was significantly lower, mainly because *P. confusa* emerges from the body of its host already at the 4$^{th}$ instar. Moreover, a larger proportion of *P. confusa* emerged from 4$^{th}$ instar larvae in *A. io* than in *A. urticae.* This difference is probably related to the difference in the development between the two butterfly species. The pupation time in *A. io* is longer than in *A. urticae* and the larvae reach a larger size, which probably explains why the parasitoid reaches maturity at an earlier larval development stage in *A. io* than in *A. urticae*.

**Table S1** showing the first and last week of occurrence of *P. confusa*, according to region and year. Weeks are expressed in numbers. In 2017, week 21 started May 22$^{nd}$, in 2018, week 20 started May 14$^{th}$.

| Year / region | 2017 | | 2018 | |
|---|---|---|---|---|
| | 1st week | Last week | 1st week | Last week |
| North | 21 | 29 | 21 | 25 |
| South | 20 | 32 | 20 | 26 |

**Supplementary Material 3: Habitat characteristics associated with *Phobocampe confusa* occurrence for buffer zone radii varying from 10 to 500m.**

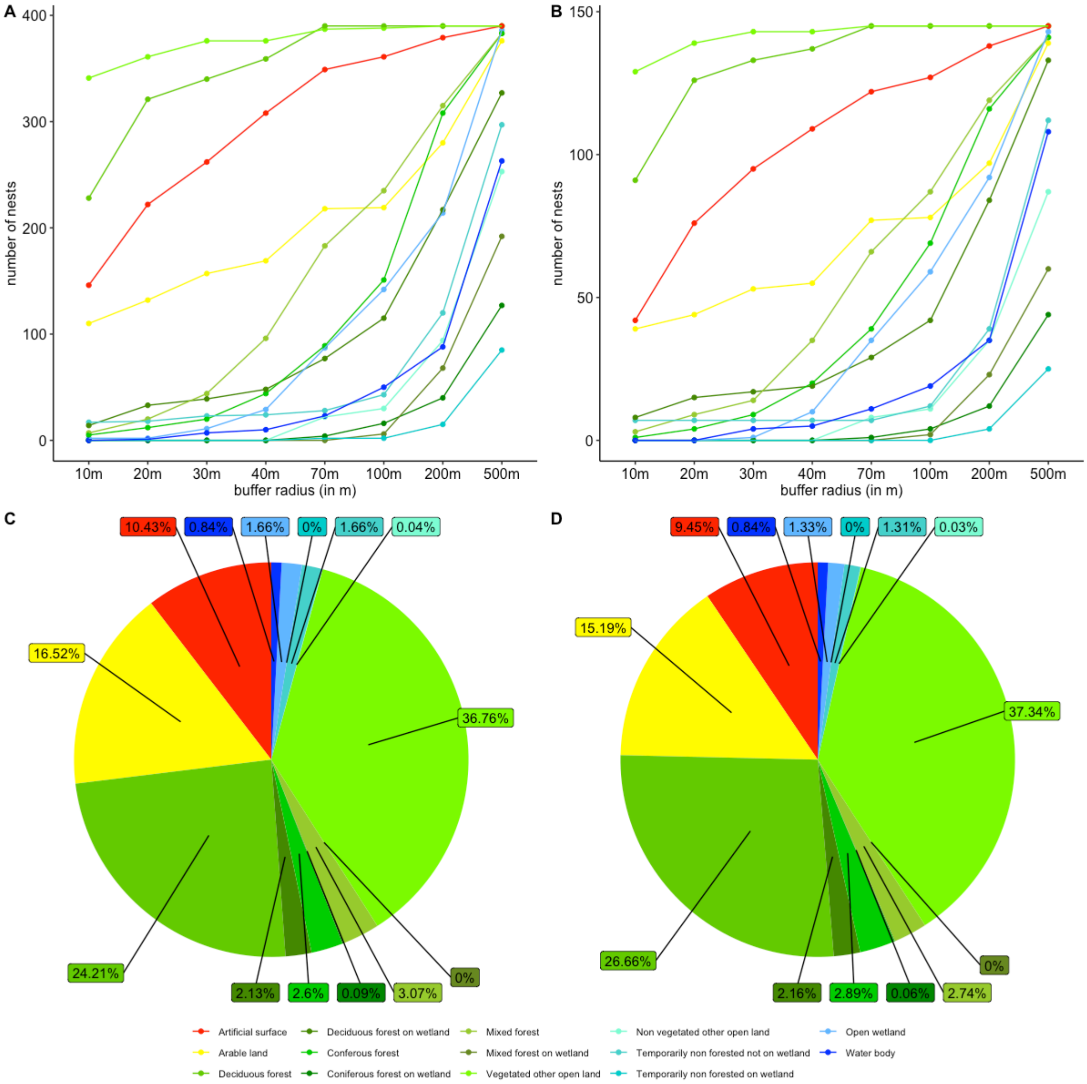


**Figure S2** Number of butterfly nests sampled surrounded by each type of land use and for each radius of the buffer zone considered (from 10m to 500m radius) for (A) all butterfly nests sampled within the phenological window of occurrence of *P. confusa* (n = 390) and (B) for the subset of butterfly nests parasitized by *P. confusa* (n = 145). In (C) and (D), pie charts representing the average land use composition within a 100m radius of the butterfly nests sampled, for all butterfly nests sampled within the phenological window of occurrence of *P. confusa* (C) and for the subset of butterfly nests parasitized by *P. confusa* (D).

**Table S2.** summary of the models built to examine the impact of the land use heterogeneity and fragmentation of the habitat on the propensity of a butterfly nest to be parasitized by *P. confusa*. We built one model per buffer zones considered (10, 20, 30, 40, 70, 100, 200 and 500m radius) in order to examine the impact of land use at different distance around each nest. The habitat variables selected in the final model are framed in red.

| Buffers size | Parameters | Intercept | Overlap | Overlap2 | 3rd instar | 4th instar | 5th instar | Artificial surface (%) | Length of edges (m) | Deciduous forest (%) |
|---|---|---|---|---|---|---|---|---|---|---|
| 10 m | estimate ± se | - 4.74 ± 0.80 | 10.61 ± 2.60 | -6.48 ± 2.28 | 0.82 ± 0.37 | 1.28 ± 0.34 | -0.10 ± 0.31 | -0.015 ± 0.007 | 0.015 ± 0.009 | - |
| n = 390 | z value | -5.91 | 4.09 | -2.23 | 2.23 | 3.80 | -0.32 | -2.07 | 1.67 | - |
| AIC = 429.3 | p | <0.001 | <0.001 | 0.004 | 0.026 | <0.001 | 0.75 | 0.039 | 0.095 | - |
| 20 m | estimate ± se | - 4.92 ± 0.84 | 10.90 ± 2.61 | -6.77 ± 2.28 | 0.83 ± 0.37 | 1.34 ± 0.34 | -0.090 ± 0.313 | -0.026 ± 0.009 | 0.008 ± 0.004 | - |
| n = 390 | z value | -5.86 | 4.18 | -2.97 | 2.22 | 3.94 | -0.29 | -2.97 | 2.06 | - |
| AIC = 425.0 | p | <0.001 | 0.003 | 0.003 | 0.026 | <0.001 | 0.77 | 0.003 | 0.004 | - |
| 30 m | estimate ± se | - 4.26 ± 0.77 | 10.24 ± 2.57 | -6.21 ± 2.25 | 0.82 ± 0.37 | 1.30 ± 0.34 | -0.113 ± 0.308 | -0.016 ± 0.010 | - | - |
| n = 390 | z value | -5.50 | 3.98 | -2.76 | 2.25 | 3.88 | -0.37 | -1.69 | - | - |
| AIC = 430.1 | p | <0.001 | <0.001 | 0.006 | 0.024 | <0.001 | 0.71 | 0.092 | - | - |
| 40 m | estimate ± se | - 4.47 ± 0.76 | 10.30 ± 2.55 | -6.25 ± 2.23 | 0.82 ± 0.36 | 1.30 ± 0.33 | -0.122 ± 0.307 | - | - | - |
| n = 390 | z value | -5.86 | 4.04 | -2.80 | 2.25 | 3.88 | -0.40 | - | - | - |
| AIC = 430.9 | p | <0.001 | <0.001 | 0.005 | 0.025 | <0.001 | 0.69 | - | - | - |
| 70 m | estimate ± se | - 4.47 ± 0.76 | 10.30 ± 2.55 | -6.25 ± 2.23 | 0.82 ± 0.36 | 1.30 ± 0.33 | -0.122 ± 0.307 | - | - | - |
| n = 390 | z value | -5.86 | 4.04 | -2.80 | 2.25 | 3.88 | -0.40 | - | - | - |
| AIC = 430.9 | p | <0.001 | <0.001 | 0.005 | 0.025 | <0.001 | 0.69 | - | - | - |
| 100 m | estimate ± se | - 4.10 ± 0.80 | 11.05 ± 2.66 | -7.06 ± 2.33 | 0.79 ± 0.37 | 1.29 ± 0.34 | -0.14 ± 0.31 | -0.047 ± 0.022 | - | - |
| n = 390 | z value | -5.13 | 4.15 | -3.03 | 2.16 | 3.82 | -0.46 | -2.17 | - | - |
| AIC = 428.1 | p | <0.001 | <0.001 | 0.005 | 0.031 | <0.001 | 0.65 | 0.030 | - | - |
| 200 m | estimate ± se | - 4.04 ± 0.79 | 10.81 ± 2.60 | -6.82 ± 2.27 | 0.79 ± 0.37 | 1.29 ± 0.34 | -0.14 ± 0.31 | -0.064 ± 0.027 | - | - |
| n= 390 | z value | -5.13 | 4.16 | -3.0 | 2.17 | 3.81 | -0.48 | -2.40 | - | - |
| AIC = 426.9 | p | <0.001 | <0.001 | 0.003 | 0.030 | <0.001 | 0.63 | 0.017 | - | - |
| 500 m | estimate ± se | - 4.94 ± 0.81 | 10.26 ± 2.54 | -6.25 ± 2.23 | 0.84 ± 0.37 | 1.32 ± 0.34 | -0.13 ± 0.31 | - | - | 0.026 ± 0.014 |
| n = 390 | z value | -6.14 | 4.04 | -2.80 | 2.30 | 3.92 | -0.42 | - | - | 1.85 |
| AIC = 429.5 | p | <0.001 | <0.001 | 0.005 | 0.022 | <0.001 | 0.68 | - | - | 0.065 |

**Table S3** Summary of the models built to examine the impact of the land use heterogeneity and fragmentation of the habitat on the intensity of parasitism, that is the proportion of larvae parasitized by *P. confusa*. We built one model per buffer zones considered (10, 20, 30, 40, 70, 100, 200 and 500m radius) in order to examine the impact of land use at different distance around each nest. The habitat variables selected in the final models are framed in red.

| Buffer size | parameters | Intercept | Year (2018) | Week | Week$^2$ | Overlap | Overlap$^2$ | *A. urticae* | 3rd instar | 4th instar | 5th instar | Region (South) | South x 2018 | Deciduous forest (%) | Artificial surface (%) | Arable land (%) |
|---|---|---|---|---|---|---|---|---|---|---|---|---|---|---|---|---|
| 10 m | estimate ± | - 4.63 ± | -0.11 ± | 0.67 ± | -0.084 | 6.30 ± | -5.29 ± | 0.39 ± | -0.32 ± | -0.076 | 0.16 ± | -0.24 ± | 0.78 ± | 0.005 ± | - | 0.006 ± |
| n = 145 | se | 0.78 | 0.33 | 0.27 | ± 0.036 | 1.84 | 1.57 | 0.18 | 0.19 | ± 0.175 | 0.21 | 0.19 | 0.33 | 0.003 | | 0.002 |
| AIC = | z value | -5.90 | -0.35 | 2.43 | -2.31 | 3.42 | -3.38 | 2.17 | -1.66 | -0.43 | 0.75 | -1.27 | 2.36 | 1.98 | - | 2.53 |
| 542.3 | p | <0.001 | 0.72 | 0.015 | 0.021 | <0.001 | <0.001 | 0.030 | 0.097 | 0.67 | 0.45 | 0.20 | 0.019 | 0.048 | - | 0.011 |
| 20 m | estimate ± | - 5.28 ± | -0.15 ± | 0.94 ± | -0.113 | 6.22 ± | -5.28 ± | 0.53 ± | - | - | - | -0.28 ± | 0.988 ± | -0.010 ± | - | - |
| n = 145 | se | 0.74 | 0.31 | 0.25 | ± 0.034 | 1.79 | 1.50 | 0.17 | | | | 0.18 | 0.319 | 0.003 | | |
| AIC = | z value | -7.15 | -0.48 | 3.77 | -3.29 | 3.48 | -3.52 | 3.14 | - | - | - | -1.60 | 3.10 | 3.33 | - | - |
| 544.8 | p | <0.001 | 0.63 | <0.001 | 0.001 | <0.001 | <0.001 | 0.002 | - | - | - | 0.11 | 0.002 | <0.001 | - | - |
| 30 m | estimate ± | - 4.62 ± | -0.35 ± | 0.67 ± | -0.082 | 6.04 ± | -5.10 ± | 0.42 ± | -0.30 ± | -0.065 | 0.16 ± | -0.30 ± | 1.07 ± | 0.010 ± | - | - |
| n = 145 | se | 0.77 | 0.32 | 0.27 | ± 0.036 | 1.78 | 1.50 | 0.18 | 0.19 | ± 0.174 | 0.21 | 0.18 | 0.32 | 0.003 | | |
| AIC = | z value | -6.02 | -1.10 | 2.46 | -2.27 | 3.39 | -3.40 | 2.31 | -1.59 | -0.37 | 0.76 | -1.68 | 3.34 | 3.15 | - | - |
| 543.52 | p | <0.001 | 0.27 | 0.014 | 0.023 | <0.001 | <0.001 | 0.021 | 0.11 | 0.71 | 0.45 | 0.09 | 0.019 | 0.002 | - | - |
| 40 m | estimate ± | - 4.80 ± | -0.36 ± | 0.69 ± | -0.084 | 5.67 ± | -4.81 ± | 0.47 ± | -0.35 ± | -0.141 | 0.11 ± | -0.35 ± | 1.14 ± | 0.015 ± | 0.015 ± | - |
| n = 145 | se | 0.77 | 0.32 | 0.28 | ± 0.037 | 1.78 | 1.50 | 0.18 | 0.18 | ± 0.177 | 0.21 | 0.18 | 0.32 | 0.004 | 0.006 | |
| AIC = | z value | -6.23 | -1.11 | 2.50 | -2.27 | 3.19 | -3.21 | 2.54 | -1.85 | -0.80 | 0.55 | -2.00 | 3.52 | 3.91 | 2.31 | - |
| 539.4 | p | <0.001 | 0.27 | 0.012 | 0.023 | 0.001 | 0.001 | 0.011 | 0.065 | 0.43 | 0.58 | 0.046 | <0.001 | <0.001 | 0.021 | - |
| 70 m | estimate ± | - 4.80 ± | -0.35 ± | 0.71 ± | -0.082 | 5.40 ± | -4.64 ± | 0.46 ± | -0.397 | -0.168 | 0.073 ± | -0.45 ± | 1.09 ± | 0.019 ± | 0.021 ± | - |
| n = 145 | se | 0.77 | 0.32 | 0.28 | ± 0.037 | 1.77 | 1.50 | 0.18 | ± 0.192 | ± 0.179 | 0.209 | 0.18 | 0.32 | 0.005 | 0.009 | |
| AIC = | z value | -6.20 | -1.08 | 2.56 | -2.24 | 3.05 | -3.10 | 2.51 | -2.06 | -0.94 | 0.35 | -2.58 | 3.38 | 3.55 | 2.32 | - |
| 542.4 | p | <0.001 | 0.28 | 0.010 | 0.025 | 0.002 | 0.002 | 0.012 | 0.039 | 0.347 | 0.73 | 0.010 | <0.001 | <0.001 | 0.021 | - |

**Table S3** (suite) Summary of the models built to examine the impact of the land use heterogeneity and fragmentation of the habitat on the intensity of parasitism, that is the proportion of larvae parasitized by *P. confusa*. We built one model per buffer zones considered (10, 20, 30, 40, 70, 100, 200 and 500m radius) in order to examine the impact of land use at different distance around each nest. The habitat variables selected in the final models are framed in red.

| Buffer size | parameters | Intercept | Year (2018) | Week | Week2 | Overlap | Overlap2 | *A. urticae* | 3rd instar | 4th instar | 5th instar | Region (South) | South x 2018 | Deciduous forest (%) | Artificial surface (%) | Arable land (%) |
|---|---|---|---|---|---|---|---|---|---|---|---|---|---|---|---|---|
| 100 m | estimate ± | - 4.46 ± | -0.36 ± | 0.66 ± | -0.075 | 4.44 ± | -3.80 ± | 0.43 ± | -0.436 | -0.170 | 0.052 ± | -0.52 ± | 1.07 ± | 0.018 ± | - | - |
| n = 145 | se | 0.76 | 0.32 | 0.28 | ± 0.037 | 1.76 | 1.49 | 0.18 | ± 0.194 | ± 0.180 | 0.221 | 0.18 | 0.32 | 0.005 | | |
| AIC = | z value | -5.88 | -1.11 | 2.36 | -2.01 | 2.52 | -2.55 | 2.38 | -2.25 | -0.95 | 0.25 | -2.92 | 3.32 | 3.39 | - | - |
| 542.7 | p | <0.001 | 0.27 | 0.018 | 0.044 | 0.012 | 0.011 | 0.017 | 0.025 | 0.344 | 0.80 | 0.004 | 0.024 | <0.001 | - | - |
| 200 m | estimate ± | - 4.46 ± | -0.18 ± | 0.82 ± | -0.11 ± | 4.48 ± | -3.60 ± | 0.47 ± | -0.42 ± | -0.182 | 0.061 ± | -0.44 ± | 0.84 ± | 0.015 ± | - | - |
| n = 144 | se | 0.77 | 0.32 | 0.29 | 0.04 | 1.76 | 1.48 | 0.18 | 0.19 | ± 0.179 | 0.211 | 0.18 | 0.32 | 0.005 | | |
| AIC = | z value | -5.80 | -0.54 | 2.78 | -2.73 | 2.54 | -2.43 | 2.59 | -2.18 | -1.02 | 0.29 | -2.48 | 2.59 | 2.77 | - | - |
| 527.6 | p | <0.001 | 0.59 | 0.005 | 0.006 | 0.011 | 0.015 | 0.010 | 0.029 | 0.31 | 0.77 | 0.013 | 0.010 | 0.006 | - | - |
| 500 m | estimate ± | - 4.34 ± | -0.29 ± | 0.75 ± | -0.094 | 5.40 ± | -4.47 ± | 0.41 ± | -0.319 | -0.0009 | 0.206 ± | -0.39 ± | 1.036 ± | - | - | - |
| n = 145 | se | 0.76 | 0.32 | 0.27 | ± 0.036 | 1.75 | 1.47 | 0.18 | ± 0.191 | ± 0.172 | 0.204 | 0.17 | 0.320 | | | |
| AIC = | z value | -5.70 | -0.90 | 2.76 | -2.61 | 3.09 | -3.04 | 2.27 | -1.68 | 0.005 | 1.01 | -2.28 | 3.24 | - | - | - |
| 551.42 | p | <0.001 | 0.37 | 0.006 | 0.009 | 0.002 | 0.002 | 0.024 | 0.094 | 0.996 | 0.312 | 0.023 | 0.001 | - | - | - |
